# Programmatic Detection of Diploid-Triploid Mixoploidy via Whole Genome Sequencing

**DOI:** 10.1101/371468

**Authors:** James M Holt, Camille L Birch, Donna M Brown, Joy D Cogan, Rizwan Hamid, Naghmeh Dorrani, Matthew R Herzog, Hane Lee, Julian Martinez, Undiagnosed Diseases Network, Katrina Dipple, Eric Vilain, John A Phillips, Elizabeth A Worthey

**Affiliations:** HudsonAlpha Institute for Biotechnology, 601 Genome Way, 35806, Huntsville, AL, USA; Vanderbilt University Medical Center, 1211 Medical Center Dr, 37232, Nashville, TN, USA; University of California Los Angeles School of Medicine 10833 Le Conte Ave, 90095, Los Angeles, CA, USA; Children’s National Health System, 111 Michigan Ave NW, 20010, Washington DC, USA

**Keywords:** mixoploidy, mosaicism, whole genome sequencing

## Abstract

**Purpose:** Mixoploidy is a type of mosaicism where an organism is a mixture of cells with different numbers of chromosomes. There are a broad range of phenotypes associated with mixoploidy that vary greatly depending on the fraction of cells that are non-diploid, their chromosome number, their distribution, and presumably the specific variation present in the patient. Clinical detection of mixoploidy is important for diagnosis.

**Methods:** We developed a method to detect mixoploidy from clinical whole genome sequencing (WGS) data through the identification of excess of variant calls centered on unusual B-allele frequencies. Our method isolates the signal from these variants using trio calls and then solves a basic linear equation to estimate levels of diploid-triploid mixoploidy within the sample.

**Results:** We show that our method reflects the results from a cytogenetic test. We provide examples detailing how our method has been used to identify diploid-triploid mixoploid individuals from within the NIH Undiagnosed Diseases Network. We present confirmatory findings obtained by clinical cytogenetic testing and show that our method can be used to identify the diploid-triploid ratio in these cases.

**Conclusion:** WGS data from patients with rare diseases can be used to identify mixoploid individuals. Individuals with certain characteristics as discussed should be tested for mixoploidy as part of standard clinical pipeline procedures. Scripts that perform this calculation are publicly available at https://github.com/HudsonAlpha/mixoviz.

## Introduction

Genetic mosaicism refers to the presence of multiple genetically distinct cell populations in an individual derived from a single fertilized egg [1]. This single egg criterion distinguishes mosaicism from the related phenomenon of chimerism, which describes an individual with multiple cell lineages resulting from fusion of two or more distinct zygotes [2]. Mosaicism can be in somatic and/or germ line cells depending upon the timing of the event during the development or lifespan of the organism [1, 3]. The mutation distribution pattern is in large part determined by normal cell replication, cell migration, and apoptosis during embryogenesis (for review see [4]). Mosaicism frequently occurs in early, but occasionally in later, embryogenesis because of selection against abnormal cells (particularly true for trisomic as opposed to aneuploid cells) or due to mitotic arrest and early wastage of abnormal embryos [4, 5, 6, 7]. Many recognizable mosaic patterns in the skin have been identified including narrow lines of Blaschko, broad lines of Blaschko, checkerboard pattern, phylloid pattern, and patchy pattern without midline separation (reviewed in [4]). Mosaicism has been reported for many types of chromosomal segments as well as whole chromosomes and structural abnormalities [8, 9]. Mosaicism for chromosomal segments (segmental mosaicism) is much less frequent (about 15% of all detected cases [9]) than mosaicism for whole chromosomes such as trisomies and aneuploidies. In some cases, it is clear that the presence of the molecular variation would be lethal in its constitutional state and can thus manifest only as a somatic disorder [4].

One type of whole chromosomal mosaicism is diploid-triploid mosaicism (DTM) or 2n/3n mosaicism, in which individuals have a percentage of triploid cells with 69 chromosomes in addition to diploid cells [10, 11, 12, 13]. In contrast to patients with full triploidy, which occurs in approximately 1% of all conceptions and almost always results in early spontaneous abortion, patients with mixoploidy can survive the neonatal period and in some cases live to adulthood [11, 12, 14].

A number of different mechanisms of origin for DTM have been suggested. The first is a meiosis I or II error, leading to an egg or sperm with two copies of the genome from a single parent followed by post-zygotic diploidization of some cells [15]. The second is delayed digyny, by incorporation of a pronucleus from a second polar body into one embryonic blastomere [16, 17]. The third is delayed dispermy, a type of diandry that occurs by incorporation of a second sperm pronucleus into one embryonic blastomere [17, 18]. The last is chimerism with karyotypes from one diploid and one triploid zygote fusing and developing into a single individual [16, 19].

It has been suggested that neither mosaicism nor chimerism were appropriate terms for DTM, based on the fact that chimerism requires the contribution of two independent zygotes and the fact that mosaicism requires that the secondary cell line be derived from a pre-existing primary cell line. Therefor, we and others use the term “mixoploidy” to describe such cases simply indicating the coexistence of multiple cell lines with different ploidy levels [10, 19, 20].

The mechanism leading to their phenotype in mixoploid individuals is poorly understood [12, 21, 22]. Abnormalities may result from an over-expression of dosage-sensitive genes or alternatively due to an imbalance in gene expression caused by imprinted genes. The latter may explain the similarity of DTM with genetic imprinting disorders such as Silver-Russell syndrome, congenital hyperinsulinism, and Beckwith-Wiedemann syndrome [23].

Diploid-triploid mosaicism can be associated with truncal obesity, body/facial asymmetry, weak muscle tone (hypotonia), delays in growth, mild differences in facial features, fusion or webbing between fingers and/or toes (syndactyly), and irregularities in the skin pigmentation. Intellectual disabilities may be present but are highly variable and can range from mild to more severe [24, 25, 26, 27]. Several clinical conditions share these age-dependent symptoms confounding the definitive diagnosis.

Cytogenetics is often used to identify individuals suspected to be mosaic due to phenotype or uneven pigmentation. Although recognition of mosaicism is relatively straightforward, detection of low-level mosaicism (<20%) can be challenging because progressively more cells must be evaluated to detect decreasing levels of mosaicism [21, 23, 25]. Microarray-based techniques began to replace cytogenetic testing in 2005 due to increased sensitivity and the ability to test many cells simultaneously without cell culture [28, 29]. ArrayCGH can be used to identify mosaicism when variant cells constituted >10% of the total cell population [9, 30]. SNP arrays are even more sensitive than aCGH for mosaicism detection, and can detect mosaicism involving <5% of cells [30]. Often only patients with pigmentation streaks and diffuse, systemic abnormalities and dysmorphologies are selected for this type of testing.

In this study, we demonstrate that whole genome sequencing (WGS) can also identify diploid-triploid mixoploidy. The method relies on a Variant Call Format file that is typically used to identify molecular variants that may contribute to an individual’s specific clinical presentation. The method can be applied to any patient with WGS data, and indeed we run it in house on all datasets being analyzed with a WGS approach, regardless of their phenotype. This gives us the ability, in an unbiased manner, to definitively diagnose patients who might never have been identified as mixoploid and thus not considered for other types of testing.

## Material and Methods

Two patients are included in this study. At the time of testing, the first patient was a 31 month old male with an atrial septal defect, hypotonia, dysmorphic features, unilateral syndactyly, and developmental delays. He had relative macrocephaly (height was 3rd percentile, weight was 23rd percentile, and head circumference was 74 percentile). He also has a history of hepatomegaly on ultrasound that resolved over time. He was able to sit without assistance at 7 months and began standing with assistance at 24 months but was not walking by 27 months. He only used two words by 26 months and 5-10 signs. He had some behavioral problems including biting his hands. He was noted to have mild hyperpigmentation that follows the Lines of Blaschko.

At the time of testing, the second patient was a 34 month old female with short stature, absence seizures, hypotonia, global developmental delays, severe failure to thrive requiring a gastric feeding tube in infancy, areas of hypo- and hyperpigmented skin lesions on her right leg, mild dysmorphic features including a triangular face, a high pain tolerance, and premature pubic hair. She had intensive work ups including chromosomal microarray, endocrinology examination, metabolic testing, and DNA methylation testing. All test results were normal. She has had failure to thrive despite adequate calorie intake by G-tube since 12 months of age.

Both patients were identified through participation in the NIH Undiagnosed Diseases Network (UDN) [31] with clinical care being provided by the Vanderbilt University Medical Center and the University of California, Los Angeles (UCLA) School of Medicine UDN clinical site, respectively. Clinical data were obtained after written informed consent, and procedures were followed in accordance with the ethical standards of the participating institutional review boards on human research and in keeping with national standards.

In the first case, WGS was performed on two samples from the proband and one from each parent. A 4-mm human skin biopsy from both the “dark” and the “light” areas were obtained from the proband for derivation of a fibroblast culture. The dissected skin biopsy pieces were transferred into tissue culture plates containing 800 *μ*l of tissue culture media, separated, adhered to the plate, and placed into a 6-well plate in the 37° C incubator. Approximately 200 *μ*l was added every 2 days to replace any evaporated media and the amount of media increased to 2 ml of complete DMEM/20%FBS at day 2 and changed every 2-3 days. Once the cells became confluent they were trypsinized and passaged twice into T75 and then T175 flasks. Once confluent, the cells were gathered and frozen. Parental samples were obtained via blood draw.

In the second case, WGS was performed on three samples from the proband and one each parent. Samples from the proband included a blood draw and fibroblasts from “light” and “dark” skin. A skin biopsy from the proband and resulting fibroblast culture for DNA isolation was performed in a similar manner as described above. Parental samples were obtained via blood draw.

The DNA was prepared from tissue or blood samples and sequenced via standard operating protocols that have been validated for use as a Laboratory-Developed Test (LDT) in our CAP/CLIA lab. In brief, automated extraction was performed using the MagNA Pure Compact system (Roche diagnostics) in 96-well plate format. DNA was evaluated for concentration (Picogreen) and integrity (agarose gel). Approximately 480 million paired-end reads, each 150 bp in length, was generated for each sample, with typical flow-cell runs lasting 3 days each. The yields averaged over 130 Gb of sequence per sample with mean coverage of 40X over the entire reference genome. Greater than 91% of the genome was covered at 20x or more, indicating uniform and deep coverage.

After sequencing, all base calling and read filtering was performed with current Illumina software. We followed the GATK best practices to align to reference (hg19) with BWA-mem [32]. The aligned sequences were then processed via GATK for base quality score recalibration, indel realignment, and duplicate removal [33]. Over a dozen quality control metrics for each genome were collected and recorded via four Picard modules as well as in-house tools. Passing quality meant that these genomes went on to batch calling of SNVs and INDELs followed by genotyping using GATK, again according to GATK Best Practices [33]. The remaining analysis is based on the resulting Variant Call Format (VCF) file.

Datasets were analyzed using an in-house developed tool that was created to detect diploid-triploid mixoploidy in WGS datasets. The model deployed within the tool assumes a mixture of diploid and triploid cells and can be described as follows. Let *A* be a genotype inherited from the father that is present in both cell types. Let *B* be a genotype inherited from the mother that is present in both cell types. Let *C* be a genotype inherited from the mother that is only present in the triploid cells. We note that regions of *B* and *C* may be the same or different across a chromosome due to recombination events. We will refer to single binary alleles as *a*,*b*, or *c* such that 0 indicates the reference allele and 1 indicates the alternate allele. Finally, let *p* represent the frequency of diploid cells (*AB*) and (1 − *p*) represent the frequency of triploid cells (*ABC*) in the sample.

If both *p* and the parental genotype calls are known, one can calculate the expected alternate allele frequency, *f*, for a single locus as 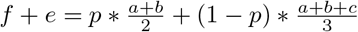 where *e* is an error term associated with the sequencing process (i.e. technical or reference bias). Note that when the parents are heterozygous, this function can lead to multiple expected allele frequencies because the inherited alleles for a single locus are unknown prior to sequencing the child. When the parents are both homozygous reference or alternate, there is only one expected frequency, *f* = 0 or *f* = 1, respectively. When they are both homozygous but for different alleles, the equation simplifies to two different functions of *p*. If the father is homozygous reference (*a* = 0) and the mother is homozygous alternate (*b* = *c* = 1), then 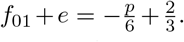 Similarly, if the father is homozygous alternate (*a* = 1) and the mother is homozygous reference (*b* = *c* = 0), then 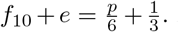. Both equations simplify to *f* = 0.56 when the sample is fully diploid with no error (*p* = 1.0, *e* = 0.0). Note that the equations above assume the source of triploidy is the mother, but the equations can be trivially modified to determine if the father is the source of the triploidy. Table 1 shows all expected frequencies for different parental genotype calls given the above assumptions

**Table 1.** Case 1 diploid-triploid expected B-allele frequencies. This table shows the expected B-allele frequencies (*f*) for a mixture of *p* diploid cells and (1 *p*) triploid cells given parental genotypes and under the assumption that the extra copy was inherited from the mother. The two corner formulas used for our estimation of triploidy are labeled as f01 and f10.

|  |  | Mother's genotype (source of triploidy) |  |  |
| --- | --- | --- | --- | --- |
|  |  | 0/0 | 0/1 | 1/1 |
| Father's genotype | 0/0 | $f = 0$ | $f = 0$<br>$f = -\frac{p}{3} + \frac{1}{3}$<br>$f = \frac{p}{6} + \frac{1}{3}$<br>$f = -\frac{p}{6} + \frac{2}{3}$ | $f_{01} = -\frac{p}{6} + \frac{2}{3}$ |
| | 0/1 | $f = 0$<br>$f = \frac{p}{6} + \frac{1}{3}$ | $f = 0$<br>$f = -\frac{p}{3} + \frac{1}{3}$<br>$f = \frac{p}{6} + \frac{1}{3}$<br>$f = -\frac{p}{6} + \frac{2}{3}$<br>$f = \frac{p}{3} + \frac{2}{3}$<br>$f = 1$ | $f = -\frac{p}{6} + \frac{2}{3}$<br>$f = 1$ |
| | 1/1 | $f_{10} = \frac{p}{6} + \frac{1}{3}$ | $f = \frac{p}{6} + \frac{1}{3}$<br>$f = -\frac{p}{6} + \frac{2}{3}$<br>$f = \frac{p}{3} + \frac{2}{3}$<br>$f = 1$ | $f = 1$ |

We then calculated the observed average and median B-allele frequencies for *f*_01_ and *f*_10_ using the data from the VCF files for our cases. We scanned variants using a strict filter that required a variant to be a bi-allelic single nucleotide variant, all variant calls in the trio to have a minimum read depth of 20 and a minimum call quality of 20, and for parental genotype calls to be homozygous for different alleles. For each variant that met this criteria, we calculated the B-allele frequency in the proband by dividing the alternate variant read depth by the total read depth to get a fraction of reads with the alternate allele. We then grouped these frequencies based on the parental genotypes and calculated both the mean and median for each category to use as the observations, *f*_01_ and *f*_10_. We calculated these values for each individual autosome and for the entire group of autosomal variants combined.

Finally, we reformed the earlier equations as a system of two linear equations where we can solve for *p* and *e*. The first equation became *p* + 6*e* = 4 − 6*f*_01_ and the second equation became −*p* + 6*e* = 2 − 6*f*_10_. This system can be trivially solved to determine the fraction of diploid and triploid cells in the population based on the observed mean and median metrics. All proband datasets were analyzed using this tool. The outputs and corresponding graphical representations were studied and compared to hundreds of other unrelated datasets from the UDN.

## Results

Unusual B-allele frequencies (Figures 1 and 3) were noted in both proband samples from case 1 and both fibroblast samples from case 2. For case 1, there were B-allele bands at frequencies other than the expected 0%, 50%, and 100% that were positioned across all chromosomes. Additionally, the pattern (but not the B-allele frequencies) was shared between the two different fibroblast samples from the proband. These unusual results were returned to the Vanderbilt clinical site with the recommendation for clinical cytogenetic testing for presumed mixoploidy.

We applied our WGS-derived B-allele frequency method to both fibroblast samples from case 1. Observed B-allele frequencies and the derived population frequencies are shown in Table 2. Using either the mean or median calculated approximately 74% diploid cells and 26% triploid cells in the “light” skin fibroblast matching and perhaps refining the observation from cytogenetics. We ran the same calculation using the “dark” skin fibroblast sample and calculated a distribution of approximately 89% diploid and 11% triploid in the second sample. The clinical site performed cytogenetic testing on the proband’s lighter skin fibroblast sample and results returned indicated a mixture of approximately 75% diploid cells and 25% triploid cells.

**Table 2.** Case 1 calculated and observed diploidy. This table shows the observed *f*_01_ and *f*_10_ mean/median values along with the predicted diploidy fraction for both proband samples from case 1 when calculated across all autosomes. Both calculations for the light sample estimate that ~74% of cells are diploid and the remaining ~26% are triploid. Cytogenetic diploidy is the reported percentage of normal diploid cells from cytogenetic tests.

| Sample | $f_{01}$ obs | $f_{10}$ obs | Error ($e$) | Diploidy fraction ($p$) | Cytogenetic Diploidy |
| --- | --- | --- | --- | --- | --- |
| Light-mean | 0.5413 | 0.4543 | 0.0022 | 0.7389 | $\sim 0.75$ |
| Light-median | 0.5424 | 0.4545 | 0.0015 | 0.7365 | $\sim 0.75$ |
| Dark-mean | 0.5160 | 0.4787 | 0.0027 | 0.8880 | — |
| Dark-median | 0.5179 | 0.4792 | 0.0015 | 0.8839 | — |

**Figure 1.**
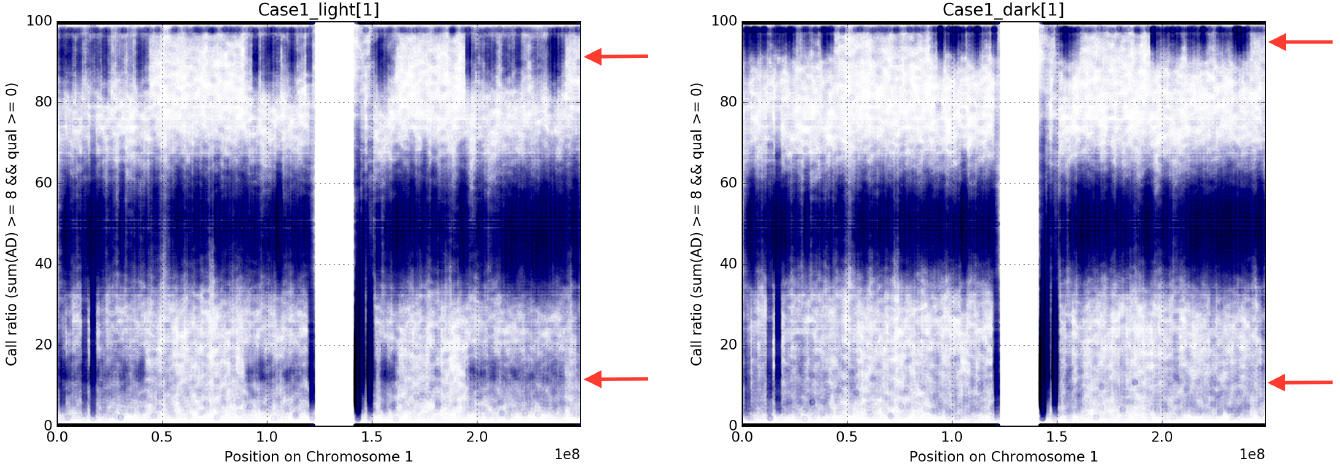
Case 1 B-allele plots. These figures show the expected diploid B-allele frequency bands at 0%, 50%, and 100% for the two fibroblast samples from case 1, light skin on the left and dark skin on the right. In addition, partial bands that do not span the length of the chromosome (see arrows), can be seen. These partial bands are present in the same regions of the chromosome in each sample.

**Figure 2.**
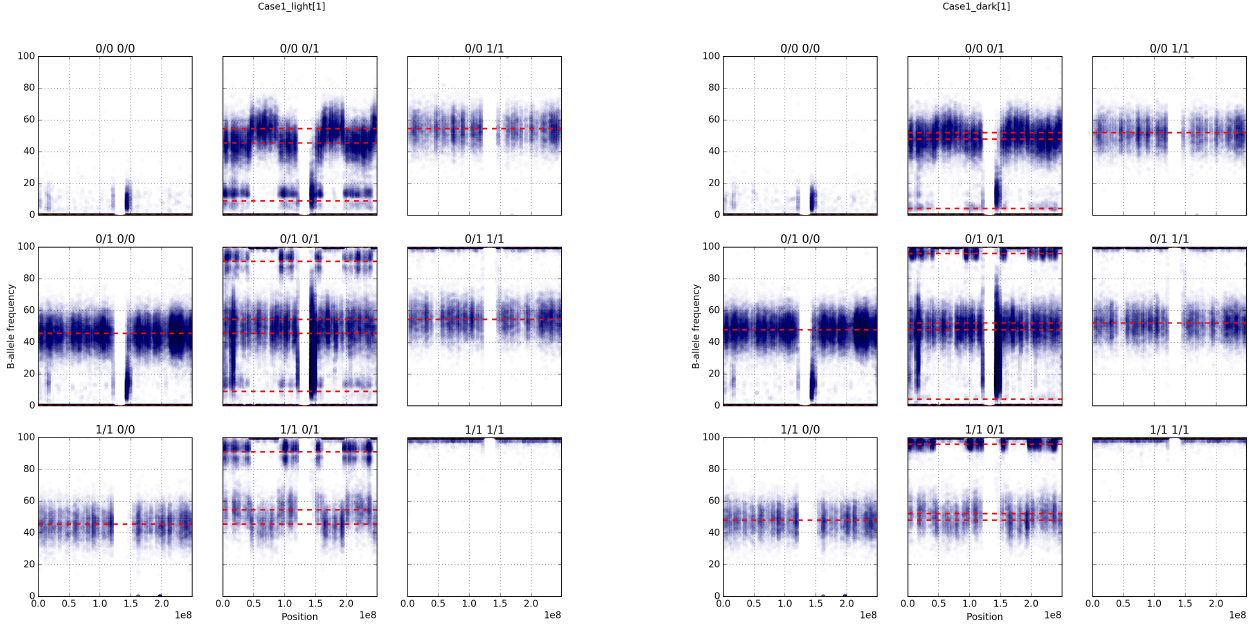
Case 1 Deconvoluted B-allele plots. These images show the B-allele frequencies for the light skin (left) and dark skin (right) fibroblasts from case 1 when split based on paternal and maternal genotype calls. Red dashed lines represent predicted B-allele frequencies derived from the formulae in Table 1. Note that each predicted frequency line resides roughly in the center of a band of observed frequencies.

**Figure 3.**
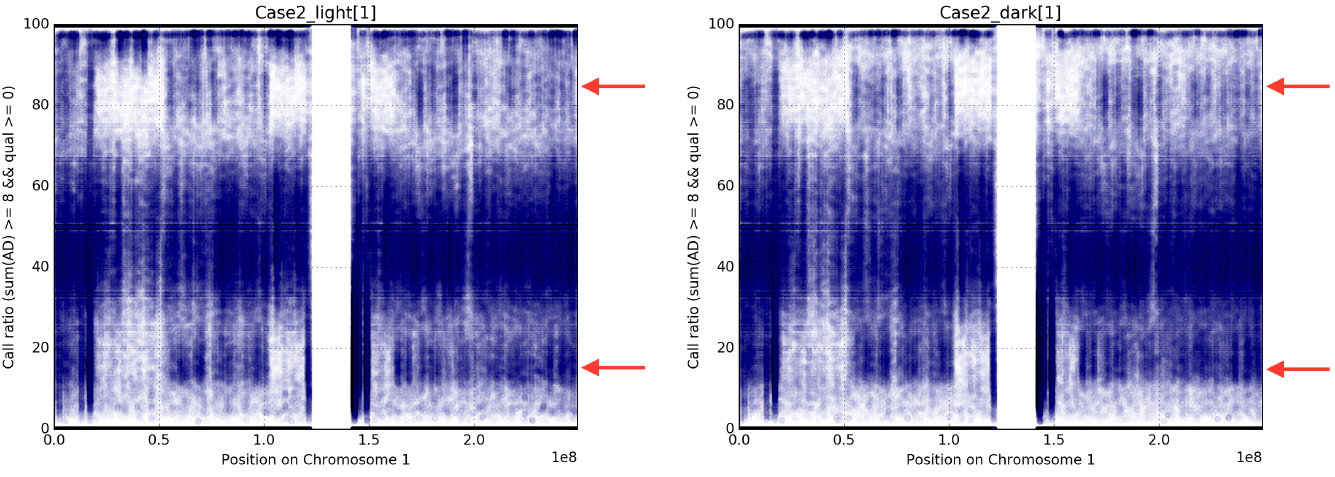
Case 2 B-allele plots. These figures show the B-allele frequencies for the two fibroblast samples from case 2, light skin on the left and dark skin on the right. Similar to case 1, there are partial bands at unusual B-allele frequencies (see arrows). Note that these partial bands cover the same regions of the chromosome in case 2, but that those regions are different than the ones from case 1.

Using the derived *p* and *e* values for each sample, we then calculated expected B-allele frequencies for all inheritance combinations. Figure 1 shows the B-allele frequency plot for chromosome 1 for both fibroblast sample where the variants have been categorized based on the parental genotype calls. All expected B-allele frequencies based on the calculated *p* have been overlaid on the figures. In general, the expected frequencies are located near the centers of observed frequency bands in the chromosome. Note that low frequency bands (*f* < 0.15) are typically difficult to observe due to artifacts in the variant calling pipeline.

We performed the same analysis for the second patient and Figure 3 shows the unusual B-allele frequencies in the two fibroblast samples. Note the presence of unusual B-allele frequencies in both fibroblasts and the partial banding of these frequencies with a different pattern than from case 1. Table 3 shows the predicted levels of diploid and triploid cells for all proband samples for case 2 (including the “normal” blood sample). Note that both fibroblast samples are estimated to have much lower levels of diploid cells and higher levels of triploid cells than those from case 1.

The results of this mixoploidy test including population frequencies were returned to the UCLA clinical site along with the recommendation to perform a cytogenetic testing for diploid-triploid mixoploidy. The orthogonal cytogenetic test confirmed diploid-triploid mixoploidy and estimated 65% diploidy (13 of 20 cells) in the “light” fibroblast sample compared to the returned 54% from the WGS-based DTM test. Scripts that perform the diploid-triploid calculation from a VCF file along with the visualizations generated in the manuscript are publicly available at 
https://github.com/HudsonAlpha/mixoviz.

**Table 3.**
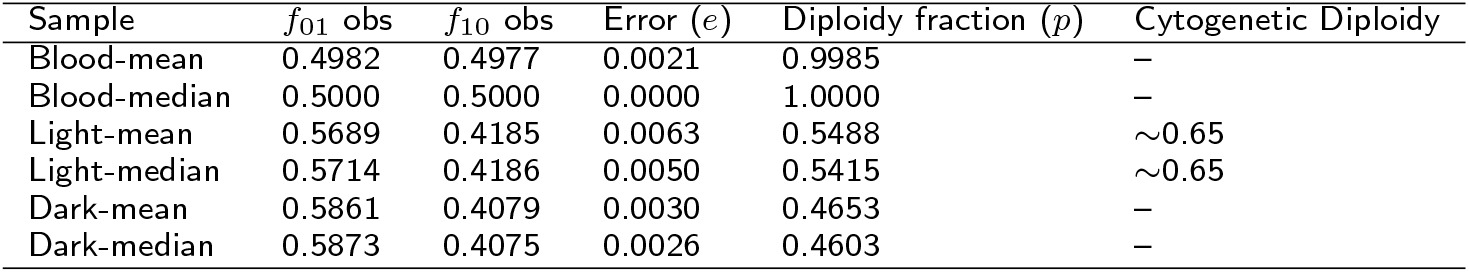
Case 2 calculated and observed diploidy. This table shows the observed *f*_01_ and *f*_10_ mean/median values along with the predicted diploidy fraction for all proband samples from case 2 when calculated across all autosomes. The blood sample is estimated to be normal (100% diploid) whereas the two fibroblast samples have relatively low levels of diploid cells (~54% in light skin and 46% in dark skin). Cytogenetic diploidy is the reported

## Discussion

We developed a method to estimate diploid-triploid mixoploidy within a WGS trio dataset. We used this method to identify mixoploidy in two of our UDN cases, both of which were confirmed through orthogonal cytogenetic testing. We then showed that our method cannot only detect mixoploidy, but can be used to obtain an estimation of the degree of mixture; similar to the results from cytogenetics. This method is extremely sensitive and, as evidenced by the different estimates in the two skin fibroblasts of both patients, it can detect when different tissues have different population mixtures. Our method is relatively simple, fast, and runs on the standard VCF file output from a WGS analysis pipeline. It can be executed as part of the WGS test reducing the need for an additional round of testing.

The output of this test can also be used to examine the specific diploid-triploid banding pattern across the patient’s genome. When combined with the WGS derived knowledge of the specific molecular variation present across the patient’s genome within those bands; we believe that this will be help us to understanding the genetic underpinnings of the patient’s specific phenotype. Work is underway to gather a significantly large dataset for this purpose.

This method is applicable to other cell type mixtures. For example, mixtures of haploid and diploid cells would yield similar formulae that can be used to estimate population frequencies in a 1n/2n mixoploidy. The method is also trivially generalizable to run on a per-chromosome basis to detect mosaic trisomies (as opposed to a full diploid-triploid mixoploid) and that test is included in our public scripts.

There are some limitations to our technique. First, it requires trio information to accurately categorize variants to perform the calculation. More complex statistical models may alleviate the need for trio information, but at least one parent would be necessary to determine the source of the extra chromosomes. Second, while some WGS analyses are tissue-agnostic, this method requires capturing the specific tissue that contains a mixture of cell types. This limitation is also true for other currently-in-use methods for detection of mosaicism.

Various mechanisms have been suggested to explain the occurrence of mixoploidy. In case 1, the partial banding pattern was seen around all centromeres, whereas with case 2 the partial banding was never seen around the centromere. This provides some information as to the possible mixoploidy mechanism in these cases. A Meiosis I error can explain either case whereas a Meiosis II error can explain the second case only. Polar body absorption (digyny) could explain either case, but is highly unlikely as it would require that the polar body happened to have opposite (or identical for case 2) centromeres for all chromosomes inherited from the mother. Dispermy or chimerism are even more unlikely due to no evidence supporting multiple haplotypes inherited from the father in either case.

## Acknowledgements

We are grateful for the participation of patients and family members and all of their referring clinicians. We would like to acknowledge all of the teams within the UDN including the coordinating center and all of the clinical sites working hard to provide definitive diagnoses for UDN patients. We would also like to acknowledge the UDN WGS core headed by Dr. Shawn Levy. This work was supported in part by the Intramural Research Program of the National Human Genome Research Institute and the NIH Common Fund through the Office of Strategic Coordination and Office of the NIH Director. Research reported in this manuscript was supported by the NIH Common Fund through the Office of Strategic Coordination and Office of the NIH Director under award numbers U01HG007530, U01HG007674, U01HG007703, U01HG007709, U01HG007672, U01HG007690, U01HG007708, U01HG007942, U01HG007943, U54NS093793, and U01TR001395. The content is solely the responsibility of the authors and does not necessarily represent the official views of the NIH.

## Competing interests

The authors declare that they have no competing interests.

